# CTLA-4 regulates PD-L1-PD-1 interactions via transendocytosis of CD80

**DOI:** 10.1101/2022.03.28.486104

**Authors:** Alan Kennedy, Max Robinson, Claudia Hinze, Erin Waters, Cayman Williams, Neil Halliday, Simon Dovedi, David M Sansom

## Abstract

CTLA-4 and PD-1 are key immune checkpoints that are targeted in the treatment of cancer. Recently it has emerged that there is a physical interaction between the ligands of these pathways (CD80 and PD-L1), which can prevent PD-L1 inhibitory functions. Since CTLA-4 captures and degrades its ligands via transendocytosis, we investigated how transendocytosis of CD80 is impacted by PD-L1 interaction. We find that transendocytosis of CD80 results in a time-dependent recovery of PD-L1 availability that correlates with CD80 removal. Moreover, CD80 transendocytosis is highly specific in that only CD80 is removed, and its heterodimeric PD-L1 partner remains on the APC. We found no evidence that CTLA-4 interactions with CD80 were inhibited by PD-L1, however efficient removal of CD80 required an intact CTLA-4 cytoplasmic domain, distinguishing this process from more general trogocytosis. We also show that simple binding of CTLA-4 to the CD80-PD-L1 heterodimer is insufficient to liberate PD-L1-PD-1 interactions, suggesting that transendocytosis of CD80 is required to effectively control PD-L1-PD-1 interactions.

## Introduction

CTLA-4 and PD-1 are two well-established immune checkpoints affecting T cell immune responses. Both pathways are involved in the regulation of autoimmunity, however deficiency in CTLA-4 and PD-1 lead to different disease outcomes. Homozygous CTLA-4 deficiency in mice leads to a fatal lympho-proliferative disease thought to be due to impaired Treg function and consequent activation of self-reactive T cells (Tivol *et al*, 1995; Walker & Sansom, 2011; Wing *et al*, 2008). Likewise in humans, heterozygous mutations have been reported to lead to a spectrum of autoimmune features (Schubert *et al*, 2014). In contrast the phenotype of PD-1 deficiency is milder but leads to exacerbated autoimmunity in several settings including a lupus-like condition in mice, thought to be related to the lack of attenuation of ongoing T cell responses (Freeman *et al*, 2000; Nishimura *et al*, 1999; Schildberg *et al*, 2016).

Despite somewhat different phenotypes, both CTLA-4 and PD-1 pathways have been targeted in cancer immunotherapy with remarkable efficacy against a range of tumours (Ribas & Wolchok, 2018; Sharma & Allison, 2015). A combination of therapies manipulating the early regulator of T cell self-reactivity (CTLA-4) and a later regulator of T cell exhaustion (PD-1) have produced outstanding results in cancer, albeit with autoimmune side effects (Larkin *et al*, 2019). Accordingly, blockade of these two pathways is attracting enormous attention and understanding their interaction is therefore important.

At the molecular level, CTLA-4 function remains incompletely understood with several proposed mechanisms (Schildberg *et al*., 2016; Walker & Sansom, 2015). However, it is clear that CTLA-4 on Treg and activated conventional T cells binds to two ligands (CD80 and CD86) found on APCs. These same ligands are used by CD28 to activate T cells and therefore interaction with higher affinity CTLA-4 reduces CD28 function (Halliday *et al*, 2020). In addition to direct competition for ligand binding, we have found that CTLA-4 is able to physically deplete its ligands from APCs in a process termed transendocytosis (Qureshi *et al*, 2011). Here, CTLA-4 binds its ligands and transfers them into the T cell followed by their degradation. This process exploits the dynamic intracellular trafficking of CTLA-4 (Qureshi *et al*, 2012) to generate a cell extrinsic control mechanism compatible with the requirement for CTLA-4 on Treg (Walker & Sansom, 2011; Wing *et al*., 2008). It is thought that this regulatory function of CTLA-4 operates constitutively in order to prevent costimulation of self-reactive T cells in the steady state, since loss of CTLA-4 function triggers profound autoimmunity in mice and humans (Kuehn *et al*, 2014; Schubert *et al*., 2014; Tivol *et al*., 1995).

In contrast to CTLA-4, the PD-1 pathway appears to control the ongoing activity of CD4 and CD8 T cells following immune activation (Wei *et al*, 2017; Wei *et al*, 2019). Like CTLA-4, PD-1 also binds to two ligands (PD-L1 and PD-L2), which whilst expressed on APCs also have a much wider tissue distribution (Schildberg *et al*., 2016). The expression of PD-1 is upregulated on T cell activation, although it is often considered to be associated with “exhausted” T cells resulting from chronic exposure to antigen either due to infection or self-reactivity (Wherry & Kurachi, 2015). Engagement of PD-1 with its ligands appears to drive this exhausted phenotype since blockade of PD-1 can reinvigorate T cells responses in viral infection and in cancer.

Despite evidence for distinct functional pathways, CD28/CTLA-4 and PD-1 pathways are connected at the molecular level. The inhibitory signalling pathway triggered by PD-1 appears to be capable of targeting both TCR and CD28 signalling. Recent studies have suggested that recruitment of the phosphatase Shp2 to PD-1 may result in a preferential targeting of the CD28 pathway over the TCR (Hui *et al*, 2017; Kamphorst *et al*, 2017), although this remains controversial (Mizuno *et al*, 2019). In addition, a direct interaction between CD80 and PD-L1 has been observed to occur with significant affinity (Butte *et al*, 2007; Butte *et al*, 2008). Whilst initial studies suggested an intercellular “*trans*” interaction between cells, more recent evidence suggests that this interaction takes place predominantly *in cis* between CD80 and PD-L1 on the same cell (Chaudhri *et al*, 2018; Sugiura *et al*, 2019). The result of this interaction is that CD80 and PD-L1 physically associate in a manner that blocks PD-L1 binding to PD-1, but potentially allows CD80 to continue its interactions with CD28 and CTLA-4. However, it remains unclear whether CD80 simply regulates the PD-1 pathway or whether PD-L1 also acts as a regulator of the CD28/CTLA-4 pathway as has been suggested (Zhao *et al*, 2019). We have therefore addressed the role of CTLA-4 transendocytosis in regulating these processes.

Given that CTLA-4 efficiently targets CD80 via transendocytosis we investigated how transendocytosis affects the PD-1 pathway. We observed that PD-L1 did not prevent CTLA-4 binding to CD80 and that CTLA-4-dependent transendocytosis remained effective despite CD80 hetero-dimerisation with PD-L1. Furthermore, despite the fact that PD-L1 was found at the immune synapse along with CD80 during transendocytosis, PD-L1 was not removed and remained on the APC. Thus CTLA-4 demonstrated a highly selective, time-dependent capacity to remove CD80 from APCs, which required the CTLA-4 cytoplasmic domain. Whilst some ligand transfer by trogocytosis was observed in cells expressing a tailless CTLA-4 mutant, this did not result in effective downregulation of CD80 from donor cells. Finally, depletion of CD80 by transendocytosis led to a time-dependent restoration of free PD-L1 across the cell membrane where it could re-engage PD-1, a feature not replicated by soluble CTLA-4 binding. These data show that CD80 is a regulator of the PD-L1: PD-1 interactions but not vice versa and that efficient ligand depletion by transendocytosis, but not simply CTLA-4 binding to CD80, is required to effectively liberate PD-L1 that can engage PD-1.

## Results

### Co-expression of CD80 and PD-L1 on the same cell disrupts PD-1 but not CTLA-4 or CD28 recognition

To study the interactions between PD-L1 and CD80 we initially used transduced cells expressing various combinations of the relevant proteins. PD-L1 was tagged with a cytoplasmic mCherry tag and probed with different anti-PD-L1 ectodomain antibodies or with CD80 Ig, PD-1 Ig or CTLA-4 Ig. When PD-L1 was expressed in isolation it was clearly detected by all PD-L1 antibodies as well as PD-1 Ig, indicating that PD-L1 was correctly expressed (**Figure 1A, expanded view 1A**). Interestingly, binding of CD80-Ig to PD-L1 was surprisingly poor, presumably reflecting the unfavourable binding of CD80 and PD-L1 in “*trans*” (Chaudhri *et al*., 2018; Sugiura *et al*., 2019). In marked contrast, co-expression of CD80 on the same cell as PD-L1 inhibited staining with several different PD-L1-specific antibodies as well as soluble PD-1 Ig (**Fig. 1B, expanded view 1A**). The level of inhibition was dependent on the antibody used, presumably reflecting different epitopes and affinities. Whilst there was a loss of PD-L1 detection upon CD80 co-expression, staining of CD80 itself was readily detectable with abatacept (CTLA-4-Ig) **(Figure 1B)**, indicating that CTLA-4 binding to CD80 was not obviously impaired by the presence of co-expressed PD-L1. Using CD28 downregulation as a measure of CD28 engagement, we also confirmed that PD-L1 had no impact on the ability of CD80 or CD86 to engage with CD28 (**expanded view 1B).** Finally, in contrast to CD80, the co-expression of CD86 had no detectable impact on the binding of PD-L1 reagents or vice versa, highlighting clear differences between CD80 and CD86 **(Fig. 1C, expanded view 1A & B).**

**Fig. 1.**
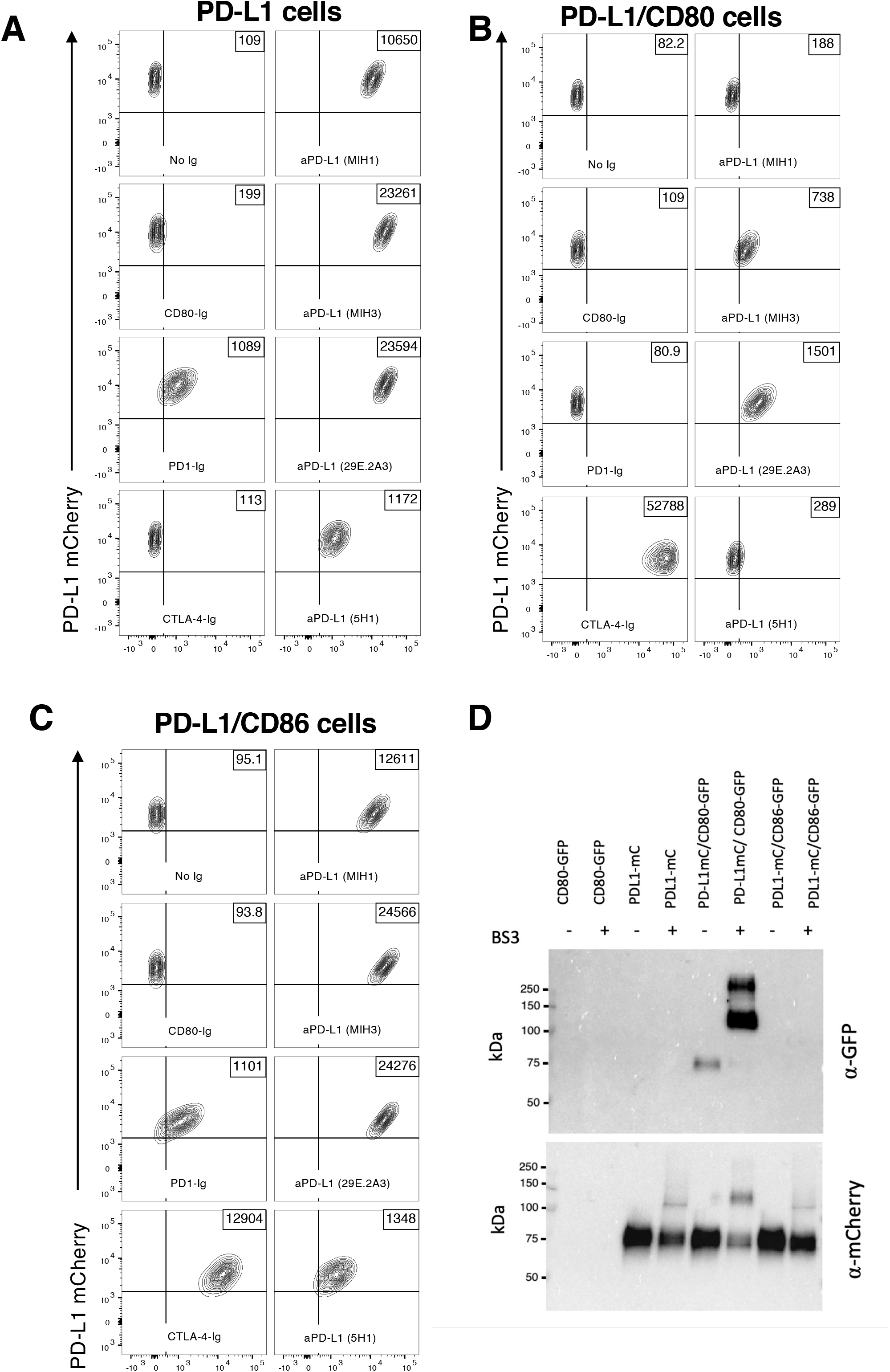
Expression of CD80 and PD-L1 *in cis* inhibits PD-L1 detection. **A-C.** CHO cells expressing PD-L1 alone (**A**) or co-expressing CD80 (**B**) or CD86 (**C**) were stained with CD80 Ig, PD-1 Ig, CTLA-4 Ig or anti-PD-L1 Ab clones MIH1, MIH3, 29E.2A3, 5H1) as indicated for 30 min at 37°C. Representative FACS plots show staining plotted against total PD-L1 mCherry levels, with MFI indicated in top right box. **D.** Western blot analysis of mCherry (PD-L1) immunoprecipitates from DG-75 cells, with and without the BS_3_ crosslinker.

Moreover, using immunoprecipitation and western blotting, we observed that PD-L1 co-precipitated CD80 in these cells upon chemical crosslinking, confirming the formation of a heterodimer between CD80 and PD-L1, which was not seen with the non-interacting CD86 **(Fig. 1D)**. Taken together these data were highly consistent with previous observations (Chaudhri *et al*., 2018; Sugiura *et al*., 2019) showing that co-expression of PD-L1 with CD80 in *cis* resulted in inhibition of PD-L1 interactions but retained CD80 binding to CTLA-4.

### Transendocytosis by CTLA-4 selectively removes CD80 but not PD-L1

Transendocytosis is a molecular process whereby the CTLA-4 ligands, CD80 and CD86, are physically removed from their host cell by a CTLA-4 expressing cell and internalised, resulting in their destruction (Qureshi *et al*., 2011). Previously it has been suggested that CD80: PD-L1 interactions can disrupt CTLA-4 binding and therefore transendocytosis (Zhao *et al*., 2019). We therefore tested the impact of PD-L1 on CD80 transendocytosis by measuring the loss of GFP-labelled CD80 ligands from donor cells as well as their acquisition by the CTLA-4^+^ recipient population using a well-established assay **(expanded view 2).** This revealed that in the presence of CTLA-4, CD80 was very effectively removed from the CTV-labelled ligand donor cells irrespective of PD-L1 co-expression (**Fig. 2A,** blue shaded quadrants). By tracking the PD-L1 mCherry signal in the same assay we observed that PD-L1mCherry remained completely associated with the donor cell despite the near total removal of CD80, indicating that CTLA-4 did not target PD-L1 for removal even when it was associated with CD80 (**Fig. 2A, Fig. 2B red quadrant)**. Thus, we concluded that whilst CD80 could prevent PD-L1 interactions with the PD-1 receptor, the reverse was not true and CD80 remained able to functionally interact with CTLA-4 and undergo transendocytosis. Moreover, the transendocytosis process demonstrated remarkable selectivity in that PD-L1 was not removed despite its CD80 partner being captured by CTLA-4.

**Fig. 2.**
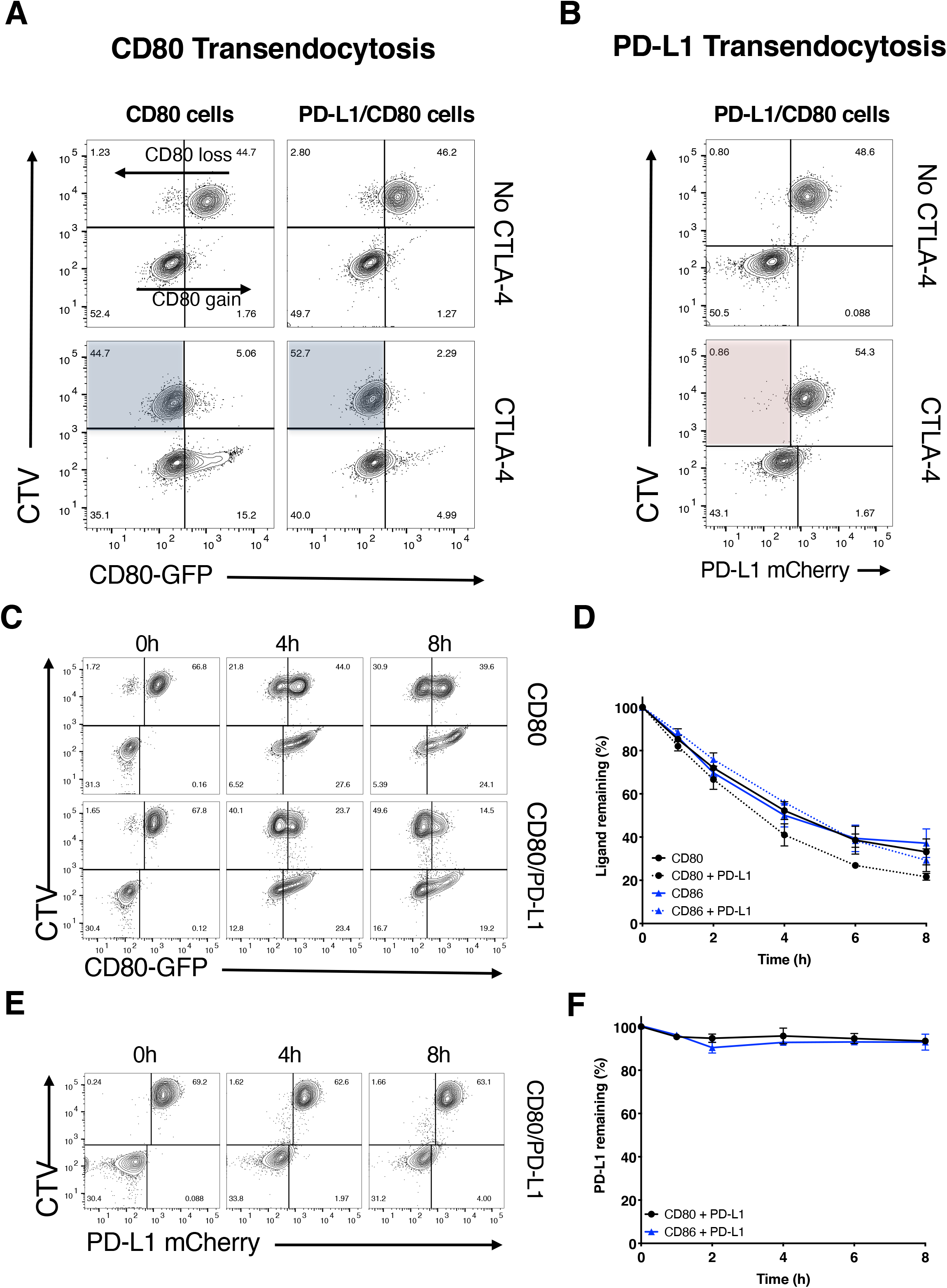
Transendocytosis of CD80 is not inhibited by PD-L1 co-expression. **A.** Transendocytosis assays were carried out overnight using CHO donor cells (CD80GFP alone or CD80GFP with PD-L1mCherry co-expression) incubated with CTLA-4 recipient cells at a ratio of 1:1. Recipient cells either lacked CTLA-4 (No CTLA-4) or expressed CTLA-4 WT. FACS plots show CD80GFP ligand loss from donor cells highlighted in blue quadrants in the presence of CTLA-4^+^ recipients. **B.** Assays as in **A** but plotted for PD-L1 expression following CD80 transendocytosis (red shaded quadrant). **C-F.** Transendocytosis assays were performed with as above but for the times indicated. Representative FACS plots showing CD80GFP expression (**C**), with full kinetic analysis of ligand downregulation on donor cells quantified in **D**. Representative FACS plots showing PDL1mCherry level at time points indicated **(E)** with full kinetic analysis of PDL1cherry level on donor cells quantified (**F**).

Given that transendocytosis is a dynamic process controlled by both the amount of CTLA-4 present and the contact time (Hou *et al*, 2015), we further monitored the impact of transendocytosis over time. As shown in **Fig. 2C,** loss of CD80 from the donor cell and its uptake by recipient cells continued over time in both the presence and absence of PD-L1. CD86 transendocytosis was also unaffected (**expanded view 3**), in keeping with its lack of interaction with PD-L1. Despite the effective removal of CD80 ligand over time **(Fig. 2D)**, there was no indication of significant PD-L1 depletion from the CD80-PD-L1 donor cells at any time point **(Fig. 2E, F)**. Taken together, the above data revealed that recognition of CD80 by CTLA-4 was not inhibited by the presence of PD-L1. Furthermore, transendocytosis of CD80 continued over extended times without removing PD-L1, demonstrating the remarkable specificity of transendocytosis and its ability to regulate PD-L1 availability.

### Efficient removal of ligand by Transendocytosis requires the CTLA-4 cytoplasmic domain

The above data showed the impact of human CTLA-4 transendocytosis expressed in CHO cells, which have proven a reliable model for transendocytosis (Hou *et al*., 2015; Khailaie *et al*, 2018; Qureshi *et al*., 2011; Schubert *et al*., 2014). To verify these observations in immune cells, we repeated these experiments using CTLA-4^+^ Jurkat T cells and B cells (DG-75) expressing matched levels of CD80 or PD-L1 **(expanded view 4)**, to allow the formation of an immune synapse. Using this system, we also investigated the role of the CTLA-4 cytoplasmic domain, since it has been suggested that CTLA-4 can effectively remove CD80 from CD80: PD-L1 co-expressing cells via trogocytosis (Tekguc *et al*, 2021), a process not requiring its cytoplasmic domain.

In line with our observations in CHO cells, CD80 was effectively downregulated over time by wild-type (WT) CTLA-4 **(Fig. 3A).** The downregulation of CD80 and CD86 was kinetically similar **(Fig. 3, A-D)** with ligand donor cells being depleted by approximately 50% after 4h. In contrast, ligand depletion by CTLA-4 Del36 (which lacks the 36 amino acids comprising the cytoplasmic domain) was much more limited, reaching only ~20% downregulation by 24h. Although the amount of ligand acquired by the CTLA-4 recipient cells appeared similar between WT and Del36 (e.g., lower right quadrants, **Fig. 3A**) this did not reflect the level of ligand downregulation from the donor cells (upper left quadrants), since ligand is continually degraded inside CTLA-4 cells. In contrast, monitoring ligand downregulation (**Fig. 3A, B -** upper left quadrants) showed that transendocytosis by WT CTLA-4 was much more effective in removing CD80 and CD86, than was trogocytosis by the Del36 mutant. This result was more notable since Del36 cells also expressed higher levels of CTLA-4 at the cell surface, due to defective CTLA-4 endocytosis **(Fig. 3E)**. We also evaluated the downregulation of PD-L1 in these experiments and confirmed again that PD-L1 remained stably associated with the donor cell in all settings **(Fig. 3C, D)**. Together these data clearly established that the CTLA-4 cytoplasmic domain was required for effective time-dependent depletion of CD80 and CD86 ligands but that PD-L1 was not effectively removed from donor cells via either transendocytosis or trogocytosis.

**Fig. 3.**
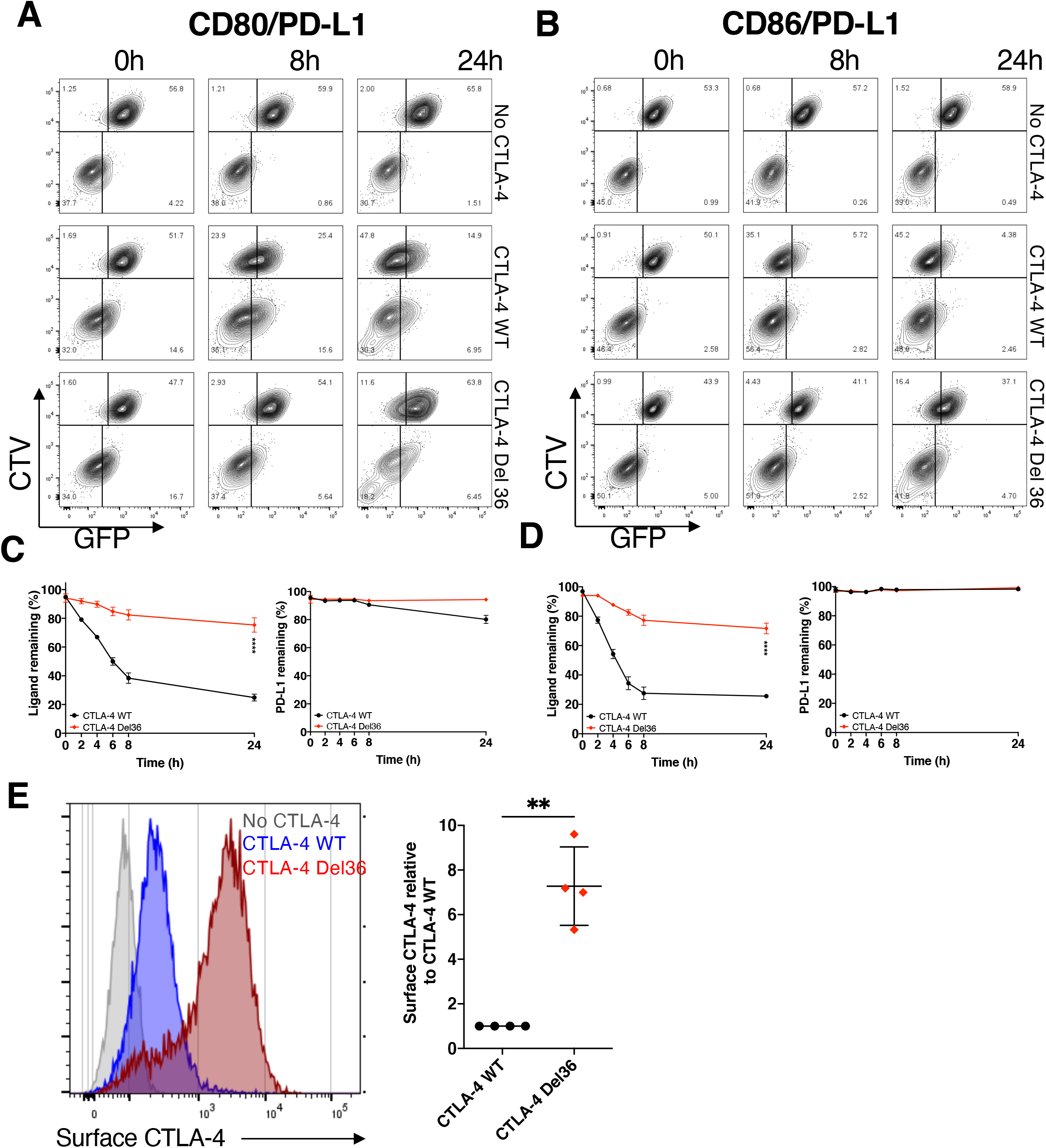
The CTLA-4 cytoplasmic domain is required for efficient transendocytosis of CD80 and CD86. **A-B.** CTV-labelled B cells (DG-75) co-expressing PD-L1mCherry and CD80GFP (**A**) or PD-L1mCherry and CD86GFP (**B**) were incubated with Jurkat cells expressing no CTLA-4, CTLA-4 WT, or a mutant CTLA-4 (Del36) lacking the cytoplasmic tail. Transendocytosis was carried out for the indicated times and analysed by flow cytometry. FACS plots show GFP ligand loss from donor B cells (upper left quadrants) and acquisition by Jurkat recipients (lower right quadrants). **C.** Levels of CD80GFP and PD-L1mCherry remaining over a 24 h transendocytosis period with CTLA-4 WT or Del36, plotted relative to no CTLA-4 control (mean ± SEM, 6 independent experiments). ****P≤0.0001, RM one-way ANOVA **D.** CD86-GFP and PD-L1mCherry remaining on donor cells over a 24 h transendocytosis period with CTLA-4 WT or Del36 were quantified and plotted relative to no CTLA-4 control (mean ± SEM, 6 independent experiments). ****P≤0.0001, RM one-way ANOVA **E.** comparison of CTLA-4 surface expression levels in Jurkat cells expressing CTLA-4 WT or CTLA-4 Del36 as determined by anti-CTLA-4 stain on ice and analysed by flow cytometry. Bar chart shows mean ± SEM from 3 independent experiments, **P≤0.01, Student’s two-tailed independent samples t-test.

### Transendocytosis of CD80 effectively liberates PD-L1 over time

Given that CD80: PD-L1 interactions *in cis* prevent the binding of PD-1, it is predicted that the physical removal of CD80 via transendocytosis should restore the availability of free PD-L1. We therefore investigated recovery of PD-1 Ig binding over time in transendocytosis assays. We observed that PD-1 Ig did not bind to cells co-expressing CD80 and PD-L1 at the outset or in the absence of CTLA-4, whereas binding to CD86/PD-L1 cells was unimpaired (**Fig. 4A & B,** top rows). However, in the presence of WT CTLA-4 (middle rows), the loss of CD80-GFP over time via transendocytosis now allowed detection of PD-L1 by PD-1 Ig. Depletion of CD80 was both CTLA-4 dependent and time-dependent, with a clear inverse correlation between the level of CD80-GFP remaining and the detection of PD-L1 at the cell surface **(Fig. 4A)**. Once again, the ability of CTLA-4 Del36 to recover PD-1 Ig binding was substantially less (albeit detectable) compared to WT CTLA-4, in keeping with its weaker ability to effectively remove CD80 **(Fig. 4A and Fig. 3A)**. As expected, whilst transendocytosis also led to reduced expression of CD86, this had no impact on PD-L1 detection, in keeping with the lack of CD86: PD-L1 interaction **(Fig. 4B)**. These data demonstrated that binding of PD-1 Ig was remarkably sensitive to changes in CD80 levels and that PD-L1 availability was related to the level of CD80 expression.

**Fig. 4.**
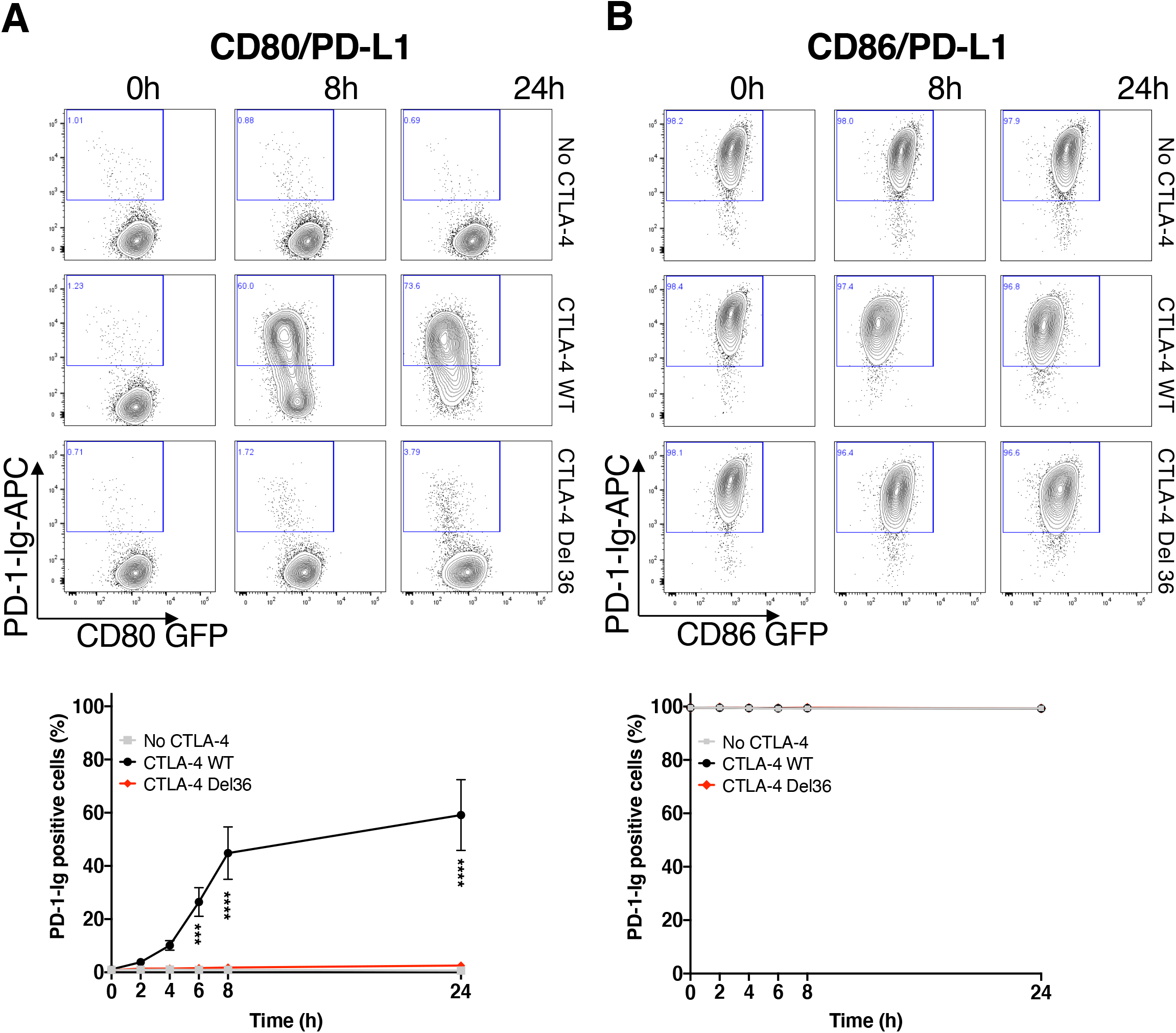
Efficient removal of CD80 by transendocytosis is required for liberation of free PD-L1 and restoration of PD-1 Ig binding. APC-labelled PD-1 Ig was used to detect PD-L1 in DG-75 cells co-expressing CD80-GFP **(A)** or CD86-GFP **(B).** Staining was carried out following transendocytosis with Jurkat cells expressing no CTLA-4, CTLA-4 WT or CTLA-4 Del36. **A** shows representative **FACS** plots of CD80-GFP vs. PD-1 Ig at the time points indicated with full kinetic data plotted below (mean ± SEM, 3 independent experiments, ***P≤0.001, ****P≤0.0001: two-way ANOVA with Tukey’s multiple comparisons test). **B.** As for A except using CD86-PD-L1 expressing cells.

We repeated these experiments using anti-PD-L1 antibodies rather than PD-1 Ig to detect PD-L1. In this case, the reduction in anti-PD-L1 staining due to CD80 co-expression was less marked than for PD-1 Ig staining, indicating that PD-L1 detection was dependent on the reagents used. Nonetheless PD-L1 staining was still impaired in the presence of CD80 **(Fig. 5A)**. Once again transendocytosis of CD80 by WT CTLA-4 markedly increased detection such that after 24h, PD-L1 staining was comparable to the level seen in the CD86 coexpressing cells, where no significant inhibition of PD-L1 staining had occurred **(Fig. 5B & C).** In keeping with our above findings, the CTLA-4 Del36 mutant did not effectively restore PD-L1 staining - although there was a limited improvement in staining following trogocytosis for 24h, this was not found to be significant. Together these data revealed differences between the ability of PD-1-Ig and antibody to detect PD-L1 as a result of CD80 interaction and that PD-1-Ig binding was generally more sensitive to inhibition by CD80.

**Fig. 5.**
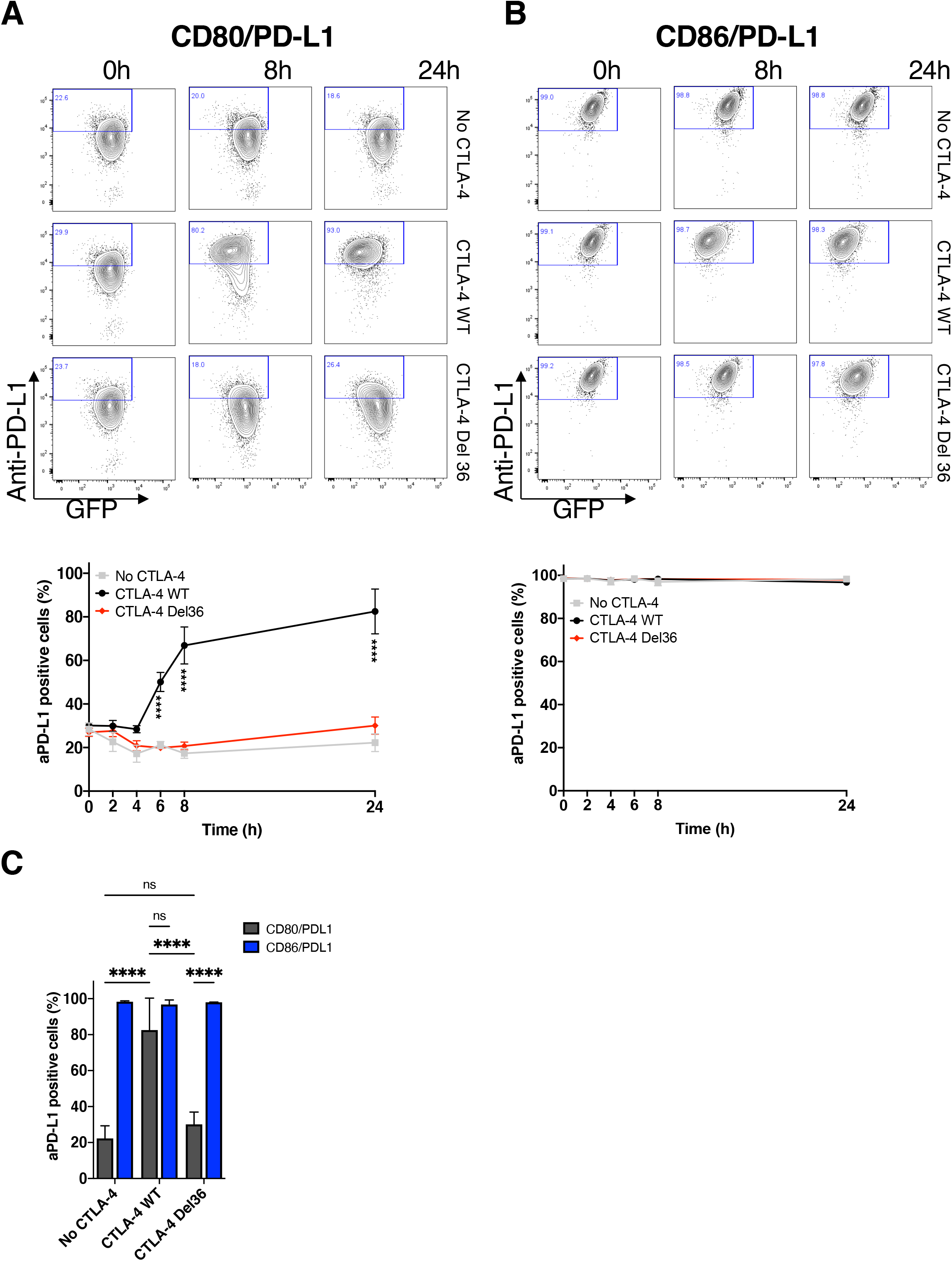
Efficient removal of CD80 by transendocytosis is required for PD-L1 detection by antibody. APC-labelled anti-PD-L1 (clone 29E.2A3) was used to detect PD-L1 in DG-75 cells co-expressing CD80-GFP **(A)** or CD86-GFP **(B).** Staining was carried out following transendocytosis with Jurkat cells expressing no CTLA-4, CTLA-4 WT or CTLA-4 Del36. **A** shows representative **FACS** plots of CD80-GFP vs. anti-PD-L1 at the time points indicated with full kinetic data plotted below (mean ± SEM, 3 independent experiments, ***P≤0.001, ****P≤0.0001: two-way ANOVA with Tukey’s multiple comparisons test). **B.** As for A except using CD86-PD-L1 expressing cells. **C.** Bar chart comparing the % anti-PD-L1 stained cells with CD80 or CD86 co-expression after 24h transendocytosis as above (mean ± SEM, 3 independent experiments, ****P≤0.0001, ns - not significant, two-way ANOVA with Tukey’s multiple comparisons test).

We also examined the time-dependent recovery of PD-1 Ig binding following transendocytosis using confocal microscopy, with similar results **(Fig. 6A)**. These data revealed clear synaptic contacts between CTLA-4^+^ Jurkat cells and B cells expressing CD80 or CD86 alongside PD-L1. In addition, transferred CD80 or CD86 could be observed as green vesicles inside of CTLA-4^+^ cells, however there was no evidence of transferred PD-L1 inside CTLA-4^+^ cells. As expected, CD86/PD-L1 cells showed clear plasma membrane staining by PD-1 Ig as shown by purple staining at all time points, consistent with CD86 having no impact on PD-1 Ig binding **(Fig. 6A** - left column). In contrast, PD-1 Ig staining gradually recovered in CD80/PD-L1 cells at later time points, as CD80 was removed by WT CTLA-4 (**Fig. 6A** - middle column). Again, the CTLA-4 Del36 mutant, whilst showing some evidence of CD80-GFP uptake, did not sufficiently deplete CD80 to enable PD-1 Ig detection **(Fig. 6A** - right column). Thus, during maintained cell-cell contacts only WT CTLA-4 could remove CD80 to reveal PD-1: PD-L1 interactions **(Fig 6B)**. Interestingly, previous data has indicated that the physical binding of CTLA-4 to the CD80-PD-L1 heterodimer, might induce dissociation of PD-L1 and CD80 (Garrett-Thomson *et al*, 2020). In such a case we might expect that PD-1 detection would focus at the synapse, where either WT or Del36 engagement of CD80 should liberate free PD-L1, even at early time points. However, we did not observe such an effect, despite clear evidence of CTLA-4: CD80 contacts (yellow synapse). Together these data highlighted a requirement for the physical removal of CD80 by CTLA-4 transendocytosis and not simple binding of CD80 by CTLA-4 in order to liberate free PD-L1 capable of binding PD-1.

**Fig. 6.**
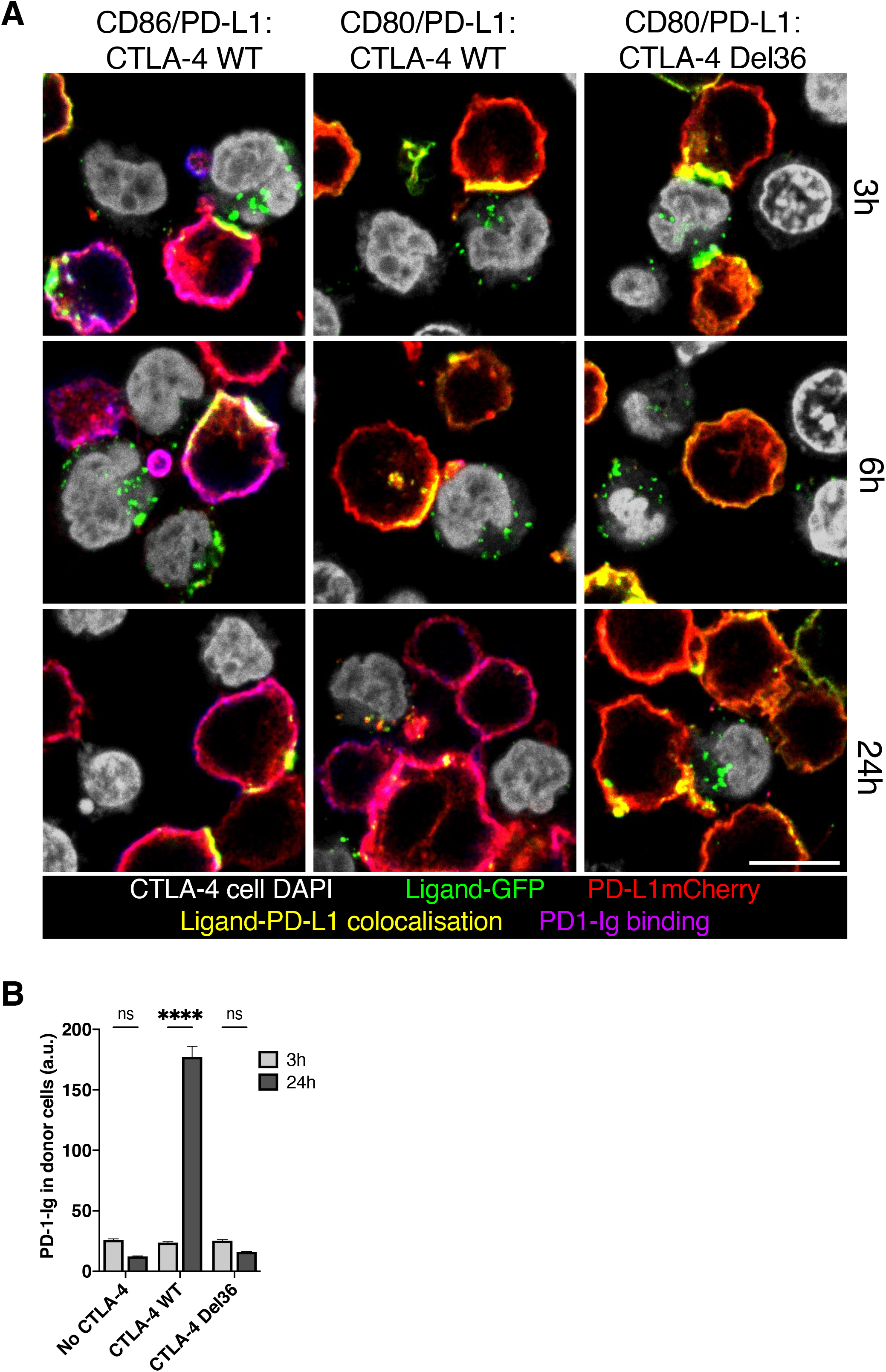
CD80/CD86 and PD-L1 are co-localised at the immune synapse during transendocytosis. **A.** DG-75 B cells expressing CD80-GFP or CD86-GFP (green) and PD-L1mCherry (red) were incubated with Jurkat cells (nuclei shown in grey) expressing CTLA-4 WT or mutant CTLA-4 Del36 for the indicated durations and analysed by confocal microscopy. 2 hours prior to assay endpoint, APC-labelled PD-1 Ig was added. CD80/CD86 and PD-L1 co-localisation at the immune synapse is shown in yellow with PD-1Ig binding shown in purple. Scale bar, 10μm. **B.** Fluorescence intensity of PD-1 Ig-APC binding to CD80/PD-L1 co-expressing B cells quantified after 3 and 24 hours transendocytosis (mean ± SEM of a minimum of 44 cells per condition, ****P≤0.0001, ns - not significant: two-way ANOVA with Tukey’s multiple comparisons test).

### Abatacept alters antibody detection of PD-L1 but does not confer PD-1 binding

The above data indicated CD80 removal by CTLA-4 transendocytosis, but not simple CTLA-4 binding, was required for liberation of competent PD-L1 expression. We therefore tested this concept in a second experiment, studying the ability of soluble CTLA-4 (abatacept) to liberate PD-L1 from CD80-PD-L1 heterodimers. It has recently been shown that soluble CTLA-4 can enhance antibody detection of PD-L1 on cells expressing both CD80 and PD-L1 (Tekguc *et al*., 2021). However, the reason for such an effect is not clear, since abatacept binds effectively to CD80 as part of a CD80: PD-L1 heterodimer and can be used to coimmunoprecipitate PD-L1 **(expanded view 5),** suggesting it does not necessarily displace PD-L1 on binding. Accordingly, using B cells expressing CD80/PD-L1 or CD80 alone, the dose response curves of abatacept binding were largely indistinguishable whether or not PD-L1 was present **(Fig. 7A and B).**

**Fig. 7.**
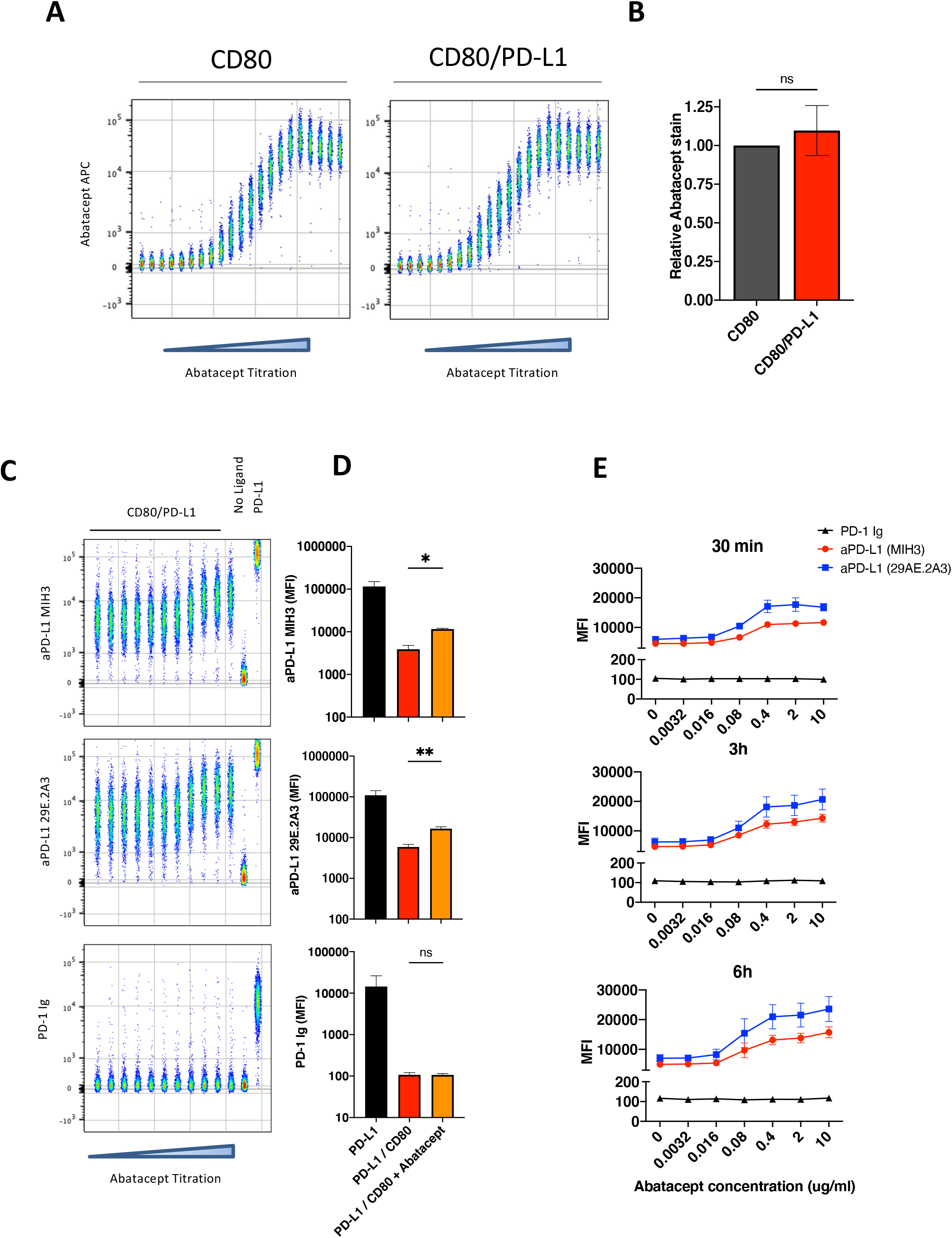
CTLA-4 Ig (Abatacept) modulates anti-PD-L1 antibody binding but fails to confer PD-1 Ig detection. **A.** Concatenated flow cytometry plots for Abatacept-APC binding to DG-75 cells expressing CD80 or PD-L1/CD80. Doses represent a serial 2-fold titration starting at 4μg/ml **B.** Relative staining of PD-L1/CD80 stained at 1μg/ml of Abatacept-APC compared to same stain on CD80 single positive cells. **C.** Concatenated plot of a 1 in 5 serial dilution of Abatacept (starting at 10μg/ml) followed by PD-L1 detection using antibody (MIH3 or 29E.2A3 clone) or using PD-1 Ig. DG-75 with no ligand or PD-L1alone (far right) are shown as staining controls in the presence of 10μg/ml of abatacept. **D.** Comparison of PD-L1 detection using PD-1-Ig or anti-PD-L1 antibodies (MIH3 or 29E.2A3 clone) with and without prior incubation with 10μg/ml of abatacept, based on data from C. **E.** Graphs showing PD-L1 detection under conditions used in C following different Abatacept incubation periods (30mins, 3h or 6h). Data is representative of 3 independent experiments showing mean ± SEM. *P≤0.05, **P≤0.01, ns - not significant.

We also studied the impact of abatacept titrations on PD-L1 detection. In keeping with the observations of Tekguc *et.al*, (Tekguc *et al*., 2021) we observed that abatacept enhanced the ability of two independent anti-PD-L1 antibodies to bind in a similar manner **(Fig. 7C and 7D)**. However, neither antibody achieved the maximal staining seen in the complete absence of CD80 or that observed following transendocytosis. Remarkably, when we attempted to detect PD-L1 with PD-1 Ig rather than antibody, we did not observe any impact of Abatacept **(Fig 7C and 7D)**. Accordingly, PD-1-Ig staining remained completely inhibited by CD80, even when abatacept was bound at high concentrations. Furthermore, increasing abatacept incubation times did not substantially increase the detection by PD-1 Ig, indicating that free PD-L1 was not released significantly over time following abatacept binding **(Fig. 7E)**. These data highlight differences between the binding of anti-PD-L1 antibodies and PD-1 Ig and indicate that release of PD-L1 due to CTLA-4 binding alone is insufficient.

## Discussion

The CTLA-4 and PD-1 pathways are critical regulators of T cell immunity in the settings of autoimmunity and cancer immunotherapy (Ribas & Wolchok, 2018; Schildberg *et al*., 2016). Understanding the roles of these distinct pathways as well as their interactions is therefore increasingly important therapeutically. It is now emerging that these two pathways are functionally connected by an interaction between the CTLA-4 ligand CD80, and the PD-1 ligand, PD-L1 when expressed on the same cell (Chaudhri *et al*., 2018) (Sugiura *et al*., 2019). Thus, it is now important to understand the impact of this interaction on both the PD-1 and CTLA-4/CD28 pathways.

Previous data on the PD-L1: CD80 interaction offers conflicting evidence as to the nature, context, and significance of this interaction. The interaction was initially reported and characterized using Fc-fusion proteins, which revealed CD80-Ig binding to PD-L1 expressing cells (Butte *et al*., 2007; Butte *et al*., 2008). Several further reports suggested that this “*trans*” intercellular interaction between ligand expressing cells resulted in inhibition of T cells responses via either CD80 or PD-L1 (Butte *et al*., 2007; Cassady *et al*, 2018; Ni *et al*, 2017; Paterson *et al*, 2011). However, more recently it has emerged that the interaction between CD80 and PD-L1 is largely precluded in the intercellular “*trans*” conformation and likely only occurs in “*cis*” where CD80 and PD-L1 are expressed on the same membrane. Alternatively interactions can occur where there is significant flexibility for binding, such as when Ig-fusion proteins are used in specific configurations (Chaudhri *et al*., 2018) (Sugiura *et al*., 2019). Our results show that CD80-Ig stained PD-L1 expressing cells very poorly, whereas in accordance with data from others we detected a robust CD80: PD-L1 *cis*interaction when both proteins are expressed on the same cell. This *cis* CD80: PD-L1 interaction precluded PD-L1 binding to PD-1 in two independent cell types (CHO and B cells). In line with the data from Sugiura *et al*. (Sugiura *et al*., 2019) we also found that CD80 forms heterodimers with PD-L1, which can be co-precipitated upon crosslinking, supporting a significant interaction between the two partners.

Structural data indicates that the heterodimeric interaction between CD80 and PD-L1 obscures the PD-1 binding site, but leaves the overlapping CD28 and CTLA-4 binding sites, which occur via the opposite (AGFCC’C”) face of CD80 (Sugiura *et al*., 2019). Interestingly the PD-1: PD-L1 interactions are unusual in structure and similar to interactions between V_H_ and V_L_ domains of an antibody, generating a “cheek to cheek” interaction (Freeman, 2008; Lin *et al*, 2008). It is the location of this binding site that appears to be obscured by CD80 when it forms a heterodimer with PD-L1. In contrast, for CD80 this interaction occurs at its normal dimerization interface, which is on the opposite face to the CD28/CTLA-4 binding site (Stamper *et al*, 2001). Accordingly, we saw no obvious effect on binding of CTLA-4 and CD28 as others have also observed (Sugiura *et al*., 2019). These data are somewhat in contrast to the suggestion of disrupted CD80: CTLA-4 interaction reported by Zhao *et al*. (Zhao *et al*., 2019). This study concluded that transendocytosis of CD80 was impaired. However, our data show continued and effective binding of CTLA-4 to CD80-PD-L1, which allows transendocytosis to proceed unimpaired, resulting in robust and continuous removal of CD80 in a time dependent manner. The reason for the differences between these observations currently remain unclear but may relate to differences in how transendocytosis was measured.

Our data further show that the outcome of CTLA-4 transendocytosis of CD80 is the liberation of PD-L1 that binds to PD-1, highlighting that CD80 is a potential regulator of the PD-1 pathway. Accordingly, in the context of PD-L1 co-expression, CTLA-4-mediated depletion of CD80 regulates PD-L1 availability, thus rescuing PD-1 receptor binding - in this context CTLA-4 becomes a regulator of PD-1 signalling. These data are consistent with those from Sugiura *et al*. (Sugiura *et al*., 2019), who showed that the level of available PD-L1 was related to the expression level of CD80 on different immune cell types. These concepts therefore have interesting implications for immunotherapy, where combinations of CTLA-4 and PD-L1 or PD-1 antibodies are used. For example, anti-CTLA-4 treatment would be predicted to increase levels of CD80 (by blocking transendocytosis), thereby potentiating CD80: PD-L1 interactions, effectively inhibiting PD-1 functions controlled by PD-L1.

Further data has suggested that CTLA-4 proteins lacking the cytoplasmic domain can effectively reveal PD-L1 after removing CD80 (Tekguc *et al*., 2021), in a process more akin to trogocytosis (Aucher *et al*, 2008; Daubeuf *et al*, 2010). Indeed, we also find that CTLA-4 when highly expressed at the cell surface (due to cytoplasmic domain deletion) is capable of trogocytosis, with CTLA-4 Del36 capturing relatively small amounts of CD80 as seen in both flow cytometric and microscopy assays. However, it is important to recognise that the tailless CTLA-4 Del36 mutant is highly over-expressed at the plasma membrane compared to WT CTLA-4, thereby facilitating trogocytosis. In our experience however, trogocytosis is frequently observed for many proteins following the physical disruption of cell conjugates that is used in order to measure protein transfer between cells using flow cytometry. In such settings one concern is that proteins get transferred as a result of this physical separation, thereby possibly overestimating the amount of protein actually captured by trogocytosis naturally. In contrast, transendocytosis robustly depletes CTLA-4 ligands *in situ*, as observed by confocal microscopy, without requiring cell separation (Qureshi *et al*., 2011). Accordingly, using confocal microscopy to analyse cells in contact, we did not observe robust detection of PD-L1 following trogocytosis. Moreover, a hallmark of transendocytosis is that it is a time sensitive process where transfer of ligands is ongoing during cell-cell contacts. This can result in the almost complete depletion of CD80 or CD86 given sufficient time. We observed here that the cytoplasmic domain of CTLA-4 was critical to this efficient time-dependent ligand transfer, with WT CTLA-4 significantly outperforming the Del36 cytoplasmic mutant. Therefore, whilst it is clear that high levels of cell surface CTLA-4 can capture significant quantities of CD80 by trogocytosis, the physiological importance of this compared to the transendocytosis process remains unclear. It is important to note that the CTLA-4 cytoplasmic domain is highly conserved through mammalian evolution, suggesting it has important conserved functions that are likely to be involved in protein trafficking (Hou *et al*, 2017; Lo *et al*, 2015; Walker & Sansom, 2011). Additionally, there are no reported mutations in the CTLA-4 cytoplasmic domain in humans despite numerous CTLA-4 mutations being reported (Schwab *et al*, 2018), attesting to the importance of this region for normal CTLA-4 function.

A further observation from our work is the remarkable selectivity of transendocytosis. Whilst trogocytosis is generally non-selective since other proteins are co-transferred during these assays (Aucher *et al*., 2008; Daubeuf *et al*., 2010; Tekguc *et al*., 2021), we found transendocytosis extremely selective in our assays. Specifically, both CD80 and CD86 were depleted from B cells and CHO cells, whilst leaving PD-L1 on the donor cell membrane. This selectivity is not cell-type dependent, or due to the nature of the cell contact, but seems due to the nature of transendocytosis itself. This is all the more remarkable for CD80: PD-L1, where despite being part of a heterodimer, only CD80 is specifically depleted, whilst PD-L1 remains. Accordingly, CD80: PD-L1 heterodimers must separate prior to transendocytosis, although this is not induced *per se* by simple CTLA-4 binding. Interestingly, CD80 binds more avidly to PD-L1 than it does to itself, with interaction affinity between CD80 and PD-L1 in the low micromolar range (Butte *et al*., 2007; Butte *et al*., 2008; Cheng *et al*, 2013). This is somewhat stronger than the interaction between CD80 monomers which form non-covalent homodimers with ~50μM affinity (Ikemizu *et al*, 2000). Nonetheless, the 0.2μM affinity between CTLA-4 and CD80 (Collins *et al*, 2002) is still highly favoured, presumably allowing CTLA-4 to specifically deplete free CD80 molecules, which may arise during normal dissociation of these non-covalent homo- and hetero-dimers.

A further significant observation from our work is that CTLA-4 binding by abatacept does not appear to dissociate the PD-L1: CD80 heterodimer effectively, such that PD-1 can bind. Our data show that soluble abatacept binding to CD80 is unimpeded by co-expression of PD-L1, consistent with the location of its binding site. Strikingly, whilst abatacept did affect staining by anti-PD-L1 antibodies, it was unable to recover detection by PD-1 Ig. Given that PD-1 Ig can readily stain free PD-L1, this suggests that abatacept does not readily generate free PD-L1. One possibility is that binding of abatacept alters the conformation of the heterodimer, such that high affinity antibodies can bind. However, in our experience this was insufficient for PD-1 binding due to affinity or epitope constraints. Given the use of abatacept clinically to treat autoimmunity and LRBA deficiency (Chitale & Moots, 2008; Lo *et al*., 2015), understanding its impact on the PD-1 pathway is of some significance. Our transendocytosis data therefore indicate that effective depletion of CD80 rather than simple binding of CTLA-4 is required to liberate functional PD-L1, although relative levels of CD80 and PD-L1 are likely to be important to the overall outcome. Such a requirement for CD80 depletion is a strong fit with the transendocytosis process itself, which is a major feature of CTLA-4 biology. Accordingly, our data suggest that liberation of PD-L1 function from CD80: PD-L1 heterodimers could not be achieved without transendocytosis.

The functional significance of the CD80: PD-L1 interaction is seen *in vivo* whereby CD80 or PD-L1 mutants lacking the ability to interact show attenuated immune responses due to excess engagement of PD-1, thereby limiting tumour immunity (Sugiura *et al*., 2019). This places CD80 as an accelerator or “turbocharger” of T cell responses by inhibiting PD-L1: PD-1 function. We would suggest that CD80 has similar functions in the CD28/CTLA-4 pathway, whereby its dominant binding to CTLA-4 could protect CD86 from transendocytosis. Together these data raise the possibility that CD80 itself operates as a switch, promoting enhanced T cell responses by inhibiting both CTLA-4 and PD-1 pathways.

## Materials and methods

### Tissue culture and cell lines

All cell lines were maintained at 37°C, 5% CO2 in a humidified atmosphere. Jurkat E6.1 T cells and DG-75 B cells were grown in complete RPMI 1640 media supplemented with 10% FBS, 2 mM L-Glutamine, 100 U/mL penicillin, and 100 mg/mL streptomycin (all from Life Technologies, Gibco).

Chinese Hamster Ovary (CHO-K1) adherent cells were maintained in complete DMEM media (Dulbecco’s Modified Eagle Medium supplemented with 10% FBS, 2 mM L-Glutamine, 100 U/mL penicillin, and 100 mg/mL streptomycin – all from Life Technologies, Gibco). Cells were routinely detached with trypsin EDTA and passaged 1 in 10.

### Cell line engineering

Transduced cell lines expressed stable integrations of transgenes using human CTLA-4, PD-L1, CD80 or CD86 tagged fusion proteins cloned into the MP71 retroviral vector. Phoenix-Amphoteric packaging cells were transfected with MP71 constructs in combination with pVSV using FUGENE HD transfection reagent (Roche Molecular Biochemical) to obtain retroviral supernatants, which were harvested 24 hrs post transfection and used to transduce CHO, Jurkat (CTLA-4) or DG-75 (CD80, CD86, PD-L1) cell lines. For transduction, non-tissue culture treated 24-well plates were coated with RetroNectin (TaKaRa) overnight at 30 mg/mL. 5×10^5^ cells were added to 1 mL of retroviral supernatants in the RetroNectin precoated wells and centrifuged at 2000 rpm at 32°C for 2 hrs. Media was changed to fresh media appropriate for each cell type 24 hrs post-infection. 3 days post-transduction, cells were stained for transduced protein expression and analysed by flow cytometry.

### Flow cytometry

#### Antibodies and detection reagents

APC conjugated PD-1-Ig (Bio-Techne, 1086-PD-050) and CTLA-4 Ig (Abatacept) were generated using the APC Conjugation Kit - Lightning-Link® according to manufacturer’s instructions (Abcam, ab201807). APC conjugated anti-PD-L1 antibodies were procured from Biolegend (clones MIH1, MIH3 and 29E.2A3).

#### Transendocytosis assays

Ligand donor cells (CHO or DG-75 B cells) expressed CD80 or CD86 molecules C-terminally tagged with GFP and/or PD-L1 molecules C-terminally tagged with mCherry. Donor cells were labelled with CellTrace Violet (CTV) labelling kit (ThermoFisher Scientific) according to the manufacturer’s instructions. CHO or Jurkat cells expressing no CTLA-4, CTLA-4 WT or Del36 were used as recipient cells. Donor and recipient cells were plated in round-bottom 96-well plates at 37°C at the ratios of donor: recipient cells and incubation times indicated in the figure legends. 20ng/ml *Staphylococcus aureus* Enterotoxin type E (Generon) was added to Jurkat: DG-75 transendocytosis assays to optimise cell-cell contacts.

Where the ability of transendocytosis of CD80 to release PD-L1 was tested, 750 ng/ml PD-1 Ig APC or 500 ng/ml 29E.2A3 were added to the transendocytosis setup 2 hrs prior to the assay endpoint. After the indicated transendocytosis duration, cells were washed three times with ice-cold PBS, fixed in 4% ice-cold paraformaldehyde in PBS for 5 min on ice and another 15 min at room temperature.

Cells were analysed by flow cytometry, gating on singlets, and GFP, mCherry or APC fluorescence measured in donor cells (CTV+).

#### Staining for flow cytometry analysis

CHO cells expressing PD-L1 alone or co-expressing CD80GFP or CD86GFP were stained with CD80-Ig, PD-1 Ig (R&D Systems), CTLA-4 Ig (Abatacept) or aPD-L1 clones MIH1, MIH3, 5h1 or 29e.2A3 at 37° C for 30 min prior to fixation in 4% PFA and permeabilisation in 0.1% saponin, and secondary stained for anti-human IgG-PE. CHO cells expressing CTLA-4 WT or CTLA-4 Del36 were surface CTLA-4 stained on ice with anti-CTLA4-PE (BD Biosciences, clone BNI3) for 60 min. Cells were washed 3 times with ice-cold PBS, fixed with ice-cold 4 % PFA in PBS for 5 min on ice and another 15 min at room temperature and analysed by flow cytometry gating on single cells.

Where the ability of soluble CTLA-4 to release PD-L1 from the CD80/PD-L1 heterodimer was tested, cells were incubated for 0.5, 3 and 6 hours with a titration of Abatacept at 37°C. Cells were washed three times and stained with either 1ug/ml PD-1 Ig-APC, MIH3 or 29E.2A3 for 30mins at 37°C. Cells were washed three times prior to analysis by flow cytometry.

### Confocal Microscopy

Transendocytosis assays were performed as described for flow cytometry at a ratio of 1:1 Jurkat: DG-75 cells. Jurkat cells expressing no CTLA-4, CTLA-4 WT or CTLA-4 Del36 were labelled with CTV and incubated with CD80-GFP, CD86-GFP and/or PD-L1mCherry expressing DG-75 cells in round-bottom 96-well plates at 37°C at a 1:1 ratio Jurkat:DG75 in the presence of 20 ng/ml *Staphylococcus aureus* Enterotoxin type E (Generon) for the indicated time points to permit transendocytosis. 750 ng/ml APC-conjugated PD-1 Ig (see flow cytometry) was added 2 hours prior to transendocytosis endpoint. Cells were then washed twice in ice-cold PBS and resuspended in cold PBS. 2×10^5^ cells were transferred into a well of a 0.01% Poly-L-lysine (Sigma Aldrich)-coated 96 well plate (Greiner Screenstar) on ice. Ice-cold 8 % paraformaldehyde in PBS was added 1:1 v/v and cells were fixed onto the well-bottom by centrifuging at room temperature for 20 min at 500g. Following sequential washes of 2% FBS in PBS, PBS and 0.1 % Saponin, cells were stained with 2μg/ml DAPI for 45min at room temperature in the dark. After sequential 0.1 % Saponin, PBS and deionised water washes, cells were mounted in Mowiol with 2.5 % DABCO.

All confocal data were acquired on an inverted Nikon Eclipse Ti equipped with a 60X oil immersion objective. Constant laser powers and acquisition parameters were maintained throughout. Digital images and scale bars were prepared using Fiji. Quantitation was performed using CellProfiler analysis software.

### Biochemistry

For crosslinking, DG-75 cells expressing CD80-GFP, PD-L1mCherry, CD80-GFP/PD-L1mCherry and CD86-GFP/PD-L1mCherry were washed three times in PBS and resuspended at 25×10^6^ cells/ml in 1mM of BS3 crosslinker (ThermoFisher, cat. Number A39266). Cells were incubated at room temperature for 30 mins followed by quenching with RPMI for 15 mins; cells were then washed three times in PBS.

Cell lysates were prepared in lysis buffer containing 50mM Tris-HCl, pH 7.4, 50mM NaCl, 10mM EDTA and 1% Triton x-100 supplemented with Protease Inhibitor Cocktail (cat. Number: 5871, Cell Signalling Technology). After 30mins incubation at 4°C with gentle rotation, lysates were spun at 10,000xG for 10 minutes. Supernatants were denatured at 100°C and processed with the Bio-Rad MiniProtean Tetra Cell Gel Electrophoresis system. Gels were transferred to PVDF membranes and blocked for 1 hour at room temperature in TBST supplemented with 5% milk. Western blotting was performed overnight at 4°C with the following antibodies: anti-GFP cat. Number: 3H9, Chromotek; anti-RFP cat. Number: 6G6, Chromotek. Blots were washed three times in TBST before secondary staining with HRP-linked anti-Mouse (Cell Signalling Technology 7076) at room temperature for 45mins. After incubation, blots were washed three times in TBST. To visualise the bands, blots were incubated for 5 minutes in Bio-Rad Clarity ECL substrate; blots were acquired on Bio-Rad Chemidoc.

For co-immunoprecipitation, supernatants from cell lysates were processed with an RFP-trap (Chromotek) or with Dynabeads protein A/protein G (Thermofisher) combined with Abatacept. Beads + lysates were incubated for 1 hour at 4°C with gentle rotation and subsequently washed three times in PBST. Immuno-precipitates were resolved by Western Blot as described for cell lysates.

## Acknowledgements

AK, EW, CH, NH and DMS were funded by the Wellcome Trust (Grants 204798, and 110297). MR was funded by an MRC industrial collaborative studentship with Astra Zeneca. CW was funded by Versus Arthritis (Grant 21147). SJD is full time employee and shareholder in AstraZeneca. MR received funding from AstraZeneca in support of this work.

This research was funded in whole, or in part, by the Wellcome Trust (Grant 204798/Z/16/Z). For the purpose of Open Access, the author has applied a CC BY public copyright license to any Author Accepted Manuscript version arising from this submission.

## Contributions

**AK** Conceptualization; Investigation; Data curation; Formal analysis; Visualization; Methodology; Writing—review and editing.

**MR;** Conceptualization;Investigation; Data curation; Formal analysis; Visualization; Methodology; Writing—review and editing.

**CH**, Conceptualization; Formal analysis; Data curation; Investigation; Visualization; Methodology; Writing—review and editing.

**EW,** Investigation; Methodology;

**NH** Investigation; Methodology;

**CW** Investigation; Methodology;

**SJD** Resources; Supervision; Funding acquisition; Writing—review and editing.

**DMS** Conceptualization; Resources; Supervision; Funding acquisition; Writing—original draft; Writing—review and editing.

## Extended view data

**EV1.**
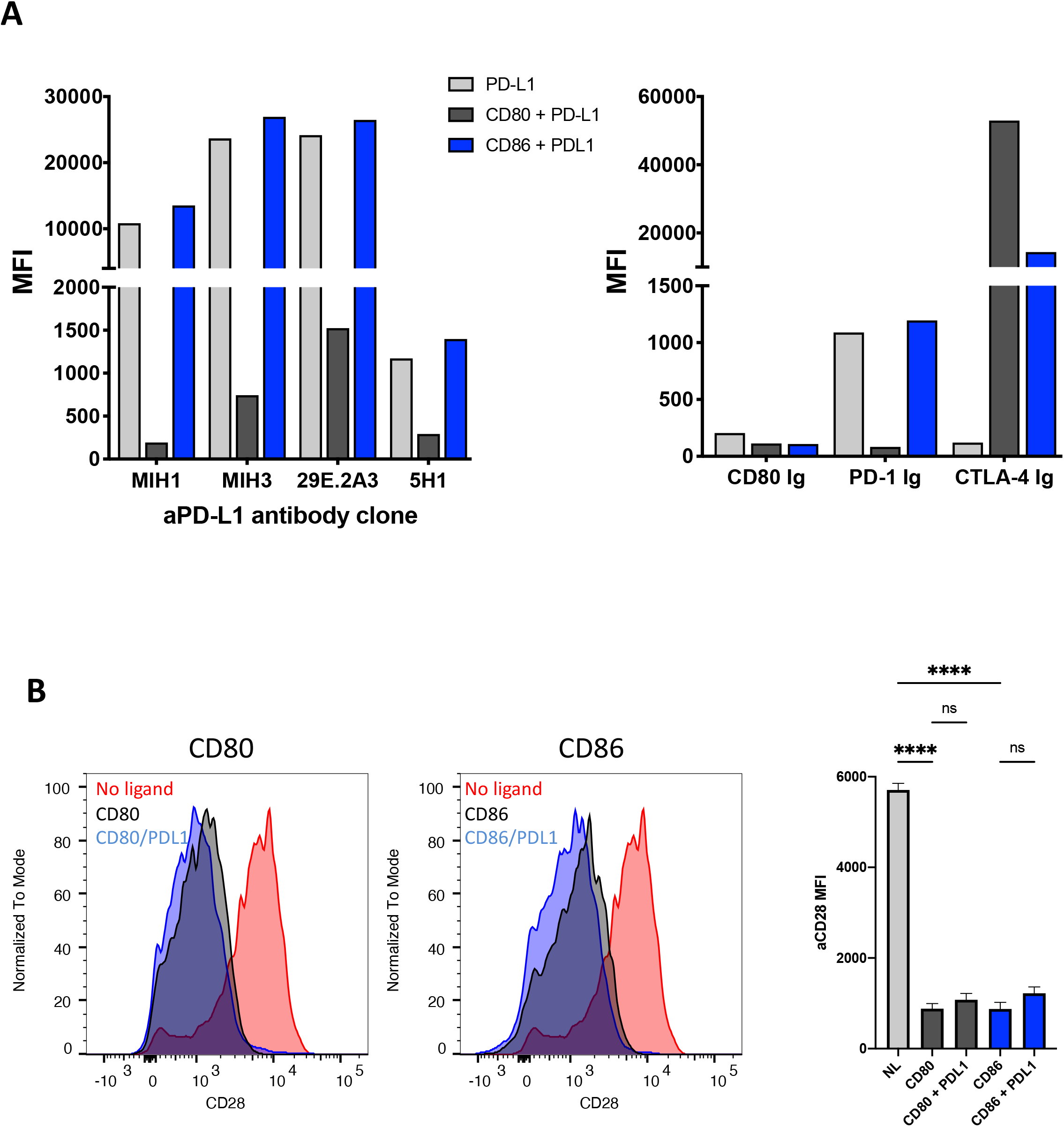
No impact of PD-L1 on detection of CD80/ CD86 by CTLA-4 or CD28. **A.** CHO cells expressing PD-L1 alone or in combination with CD80 or CD86 were stained using the reagents shown and quantified by flow cytometry. The impact of CD80 and CD86 on detection of PDL1 using anti-PD-L1 antibodies is shown (Left panel). Binding using Ig-Fusion proteins is shown (Right panel). **B.** Down regulation of CD28 following engagement by CD80 or CD86 in the presence or absence of PD-L1 is shown as a measure of CD28-ligand interaction. CD28 MFI was determined using flow cytometry is quantified in RH panel.

**EV2.**
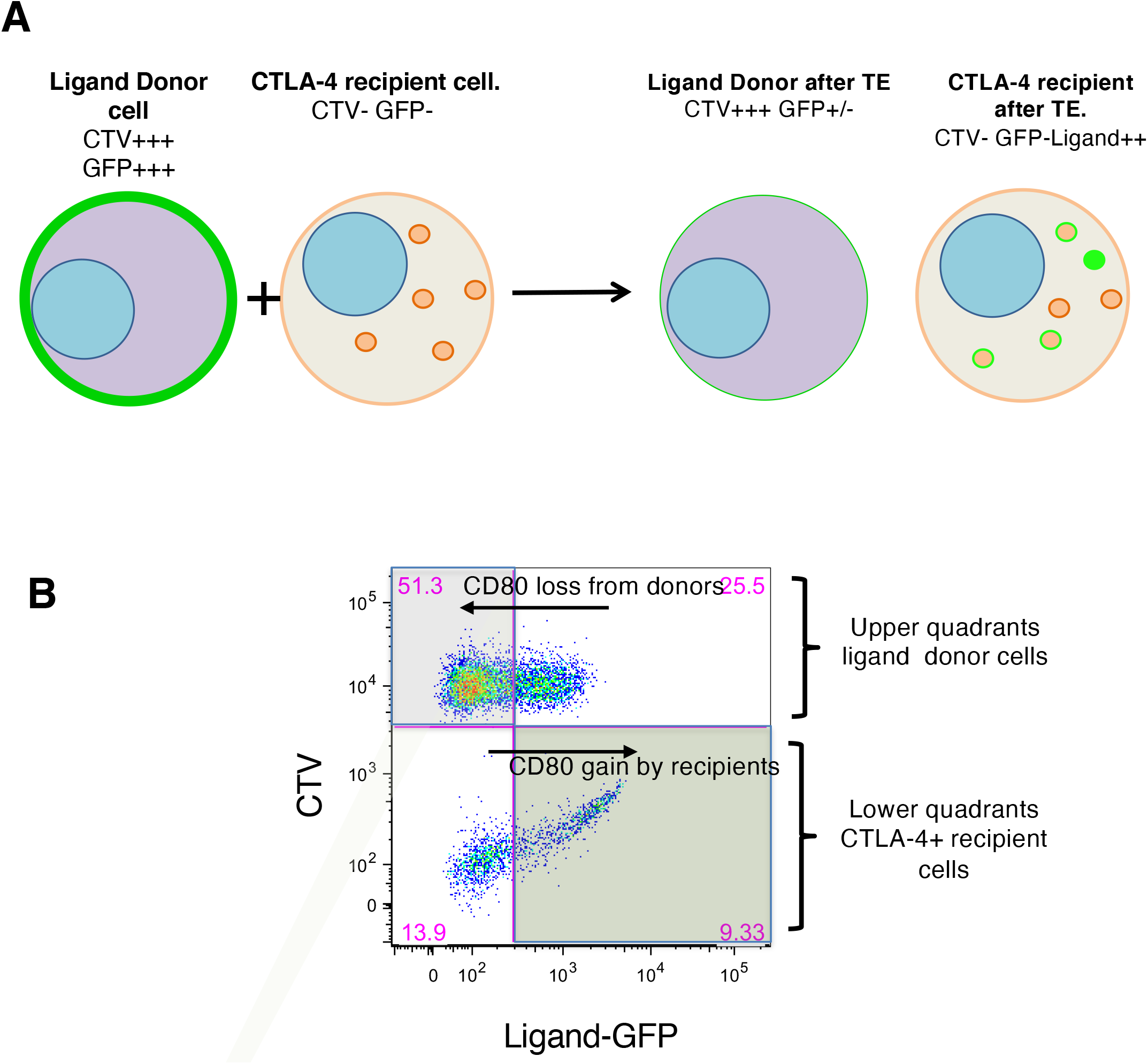
Measurement of transendocytosis by flow cytometry. **A.** Cartoon showing the principle of transendocytosis assays by flow cytometry. Ligand (donor) cells expressing CD80 or CD86 proteins with GFP fusion tags (green plasma membrane) are labelled with CellTrace Violet (CTV^+^, purple) and mixed with CTLA-4 expressing recipient cells (orange dots and membrane, CTV^-^). During transendocytosis, plasma membrane expressed ligands are removed from donor cells (reduced green plasma membrane signal) and fluorescent ligand is now detected in CTLA-4 expressing recipient cells. Internalised ligands either separate from CTLA-4 (green dots) or remain colocalized (orange dots with green outline). **B.** FACS plot shows representative result of TE, where ligand removal results in loss of GFP fluorescence from the CTV^+^ donor cell, and a concomitant increase in GFP fluorescence in the CTV^-^ recipient cells, as indicated. Note that ligand gain by recipients is subject to continuous degradation and is not a reliable indicator of total ligand transfer.

**EV3.**
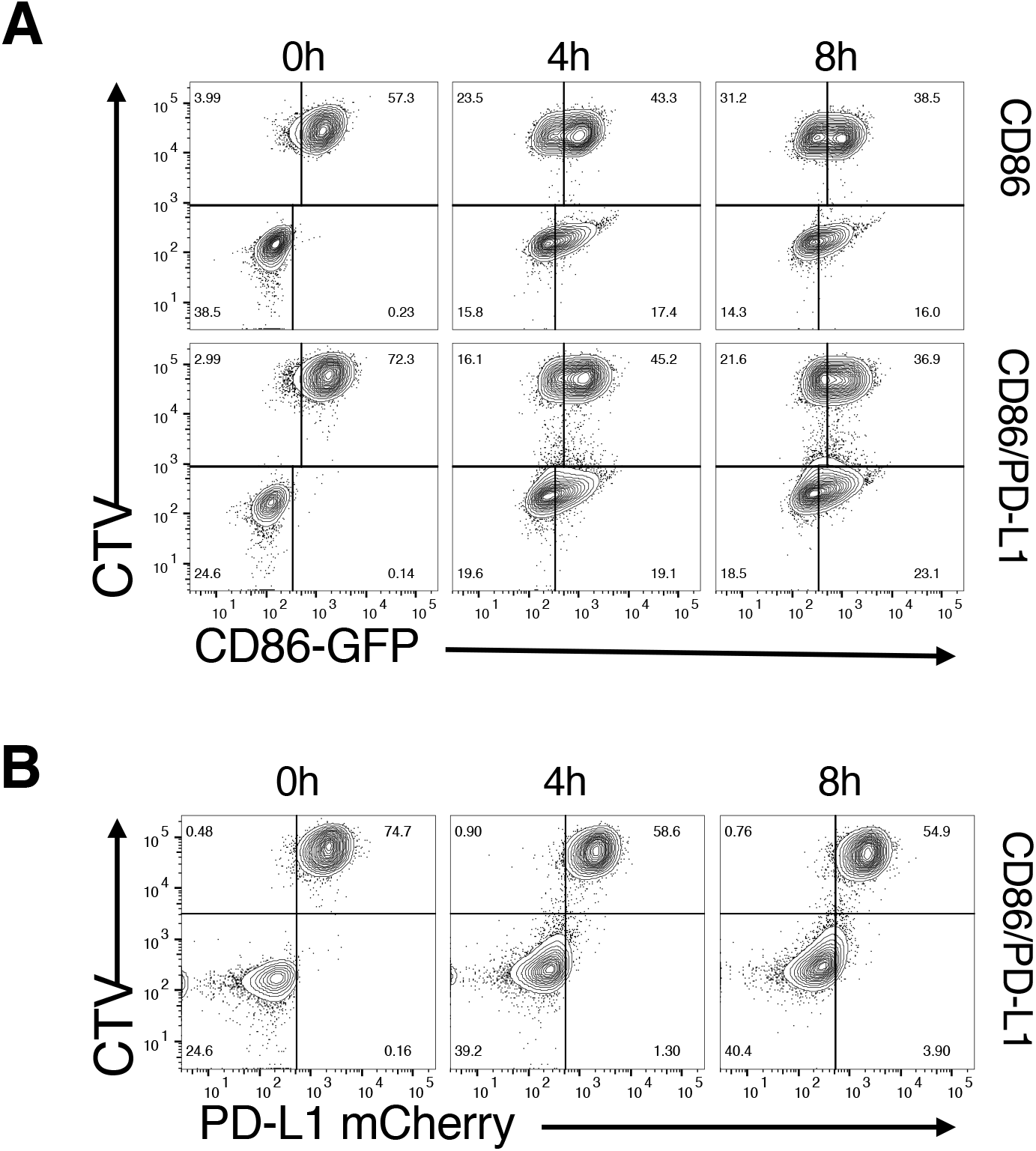
CD86 transendocytosis is unaffected by PD-L1. Transendocytosis assays were carried out using CHO cells at a ratio of 1 donor (CD86GFP alone or CD86GFP/PD-L1mCherry): 1 CTLA-4^+^ recipient. Transendocytosis assays were performed for the times indicated. Representative FACS plots show CD86GFP downregulation (**A**) and PD-L1mCherry expression (**B**) at time points indicated

**EV4.**
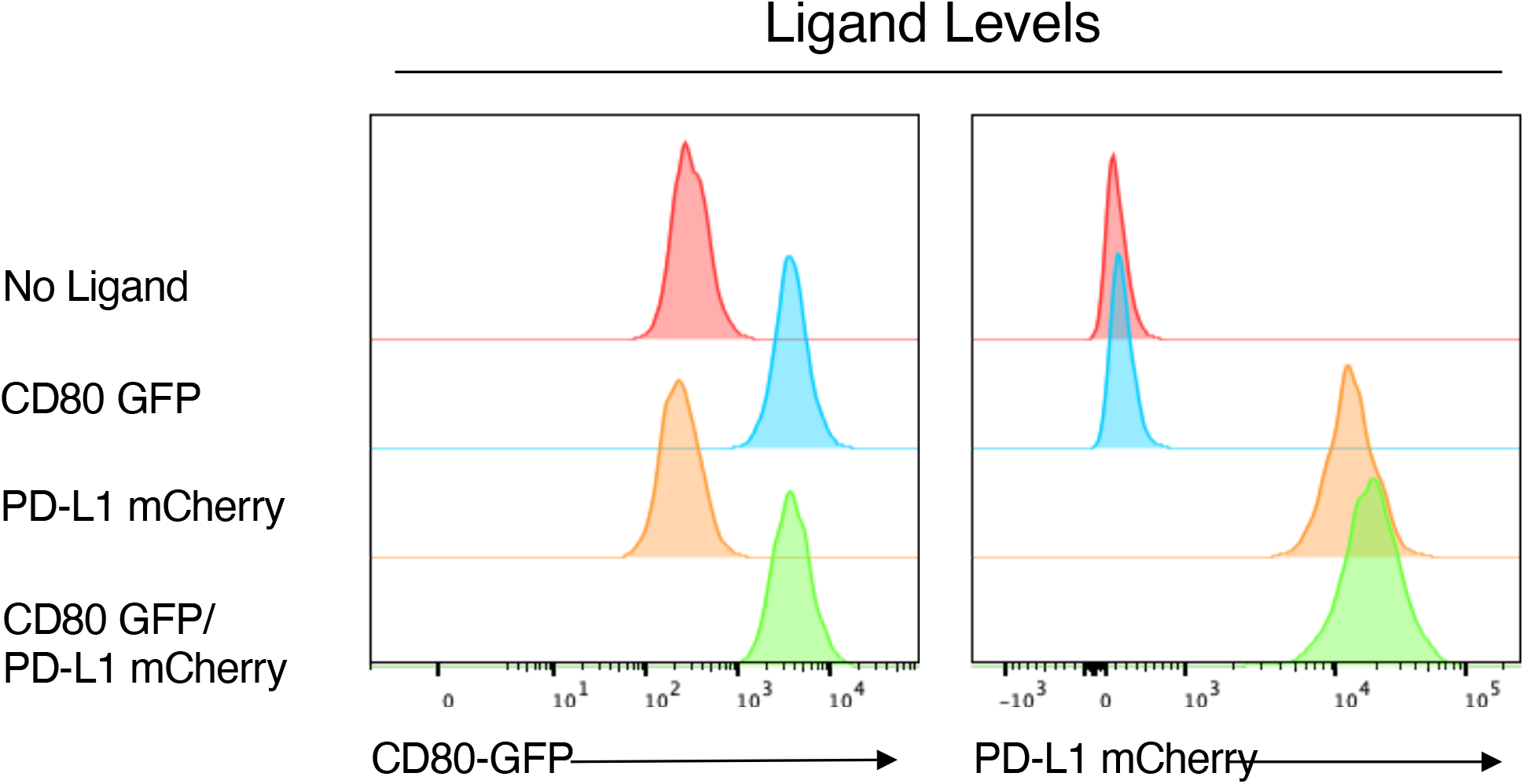
Cell lines have matched expression of ligands. Histograms showing CD80-GFP and PD-L1mCherry expression levels on transduced DG-75 cell lines.

**EV5.**
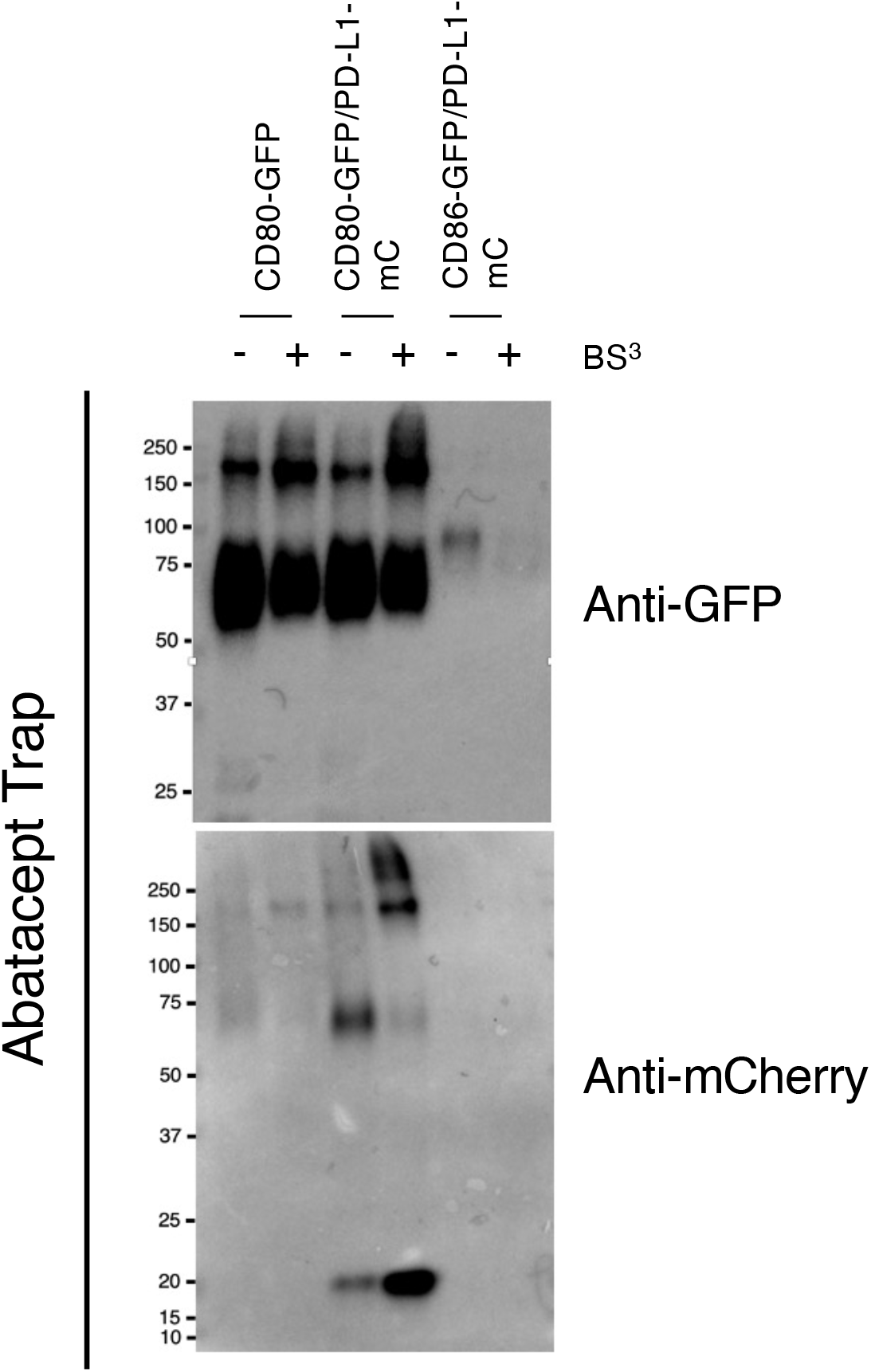
Abatacept immunoprecipitates PD-L1-CD80 interactions. Western blot analysis of Abatacept immunoprecipitates from DG-75 cells, with and without the BS_3_ crosslinker. Precipitates were immunoblotted for CD80(GFP) and PD-L1 (mCherry) as indicated.

